# Biochemical Characterization and Inhibitor Discovery for *Pf*Sir2A – New Tricks for An Old Enzyme

**DOI:** 10.1101/2024.09.25.614941

**Authors:** Dickson Donu, Emily Boyle, Alyson Curry, Yana Cen

## Abstract

The Sir2 enzyme from *Plasmodium falciparum* (*Pf*Sir2A) is essential for the antigenic variation of this parasite, and its inhibition is expected to have therapeutic effects for malaria. Selective *Pf*Sir2A inhibitors are not available yet, partially due to the fact that this enzyme demonstrates extremely weak *in vitro* deacetylase activity, making the characterization of its inhibitors rather challenging. In the current study, we report the biochemical characterization and inhibitor discovery for this enzyme. *Pf*Sir2A exhibits greater enzymatic activity in the presence of DNA for both the peptide and histone protein substrates, suggesting that nucleosomes may be the real substrates of this enzyme. Indeed, it demonstrates robust deacetylase activity against nucleosome substrates, stemming primarily from the tight binding interactions with the nucleosome. In addition to DNA/nucleosome, free fatty acids (FFAs) are also identified as endogenous *Pf*Sir2A regulators. Myristic acid, a biologically relevant FFA, shows differential regulation of the two distinct activities of *Pf*Sir2A: activates deacetylation, but inhibits defatty-acylation. The structural basis of this differential regulation was further explored. Moreover, synthetic small molecule inhibitors of *Pf*Sir2A were discovered through the screening of a library of human sirtuin regulators. The mechanism of inhibition of the lead compounds were investigated. Collectively, the mechanistic insights and inhibitors described in this study will facilitate the future development of small molecule *Pf*Sir2A inhibitors as antimalarial agents.

## INTRODUCTION

It is estimated that over one million people die annually of *P. falciparum*-related malaria with a majority under the age of five years.^1^ The development of new therapeutics against malaria is urgent due to increased resistance to current antimalarial drugs due to small changes in parasite DNA. The genome of *P. falciparum* harbors two *sir2* genes, which encode functionally similar but distinct forms of the *Pf*Sir2 protein.^2,3^ Deletion of either of these genes leads to a reduction in overall *Pf*Sir2 activity, and substantial de-silencing of a significant portion of the *var* gene family.^3,4^ Each *var* gene encodes a unique variant of the *Pf*EMP1 protein (*P. falciparum* erythrocyte membrane protein 1), the main virulence factor of *P. falciparum*. The parasite’s genome includes approximately 60 *var* genes, with all but a single variant of the *var* genes remaining transcriptionally silent, mainly due to the action of *Pf*Sir2.^2,5^ It is essential for the parasite to express only one *var* gene at a time, and to switch expression throughout the infection to evade the host’s immune response against previously expressed *var* gene products. This highly regulated gene expression is crucial for sustaining a chronic infection. Disruption of *var* gene silencing would reveal the full antigenic repertoire to the host prematurely, effectively “vaccinating” the host against all parasite variants. Evidence from analogous gene family in the intestinal parasite *Giardia lamblia* demonstrate that such silencing disruption can lead to effective vaccination in infected mice, thus supporting the potential efficacy of this approach.^6^ It is likely that *Pf*Sir2 activity will be correspondingly reduced upon inhibitor treatment and that *var* gene silencing will be disrupted.

The current study focuses on one particular *Pf*Sir2 isoform, *Pf*Sir2A. *Pf*Sir2A localizes mainly to the telomeric regions and contributes to the epigenetic regulation of antigenic variation in *P. falciparum*.^7–10^ Transcriptional profiling reveals that deletion of *Pfsir2a* induces up-regulation of a subgroup of *var* genes, particularly affecting members controlled by *upsA*, *upsC*, and *upsE* regions.^3, 4^ In line with the mutually exclusive relationship between H3K9me3 and H3K9Ac, the lack of *Pf*Sir2A activity in knock-out cells lowers the abundance of repressive H3K9me3 marks.^11^ Additionally, *Pf*Sir2A is highly enriched in the nucleolus where it fine tunes ribosomal RNA gene transcription.^12^ At the molecular level, *Pf*Sir2A exerts various biological functions through its NAD^+^-dependent protein deacetylase activity. For example, *Pf*Sir2A has been shown to remove acetyl marks from the N-terminal tails of histones H3 and H4 in the *in vitro* setting.^13, 14^ Furthermore, *P. falciparum* Alba domain-containing protein (*Pf*Alba3) has been identified as the first non-histone substrate of *Pf*Sir2A.^15^ *Pf*Sir2A-mediated deacetylation of *Pf*Alba3 at K23 enhances its DNA binding affinity, and stimulates its abasic site-driven endonuclease activity, highlighting its importance in DNA damage response.^15^ It is discovered recently that *Pf*Sir2A is also a lysine defatty-acylase.^16^ It prefers to remove medium to long chain fatty acyl groups from lysine residues, although the physiological significance of this novel activity remains elusive.

The collective biological insight into *Pf*Sir2A function suggests that the enzymatic activity is linked to the maintenance of heterochromatin, in sub-telomeric regions in particular. Multiple studies show, however, that *Pf*Sir2A deacetylates histone H3 lysine 9 (H3K9Ac) peptides with a slow turnover rate, several orders of magnitude slower than other sirtuins.^14, 16^ The weak *in vitro* activity cannot reconcile with the biological observations, suggesting that peptide-independent mechanisms promote more robust *Pf*Sir2A activity. In the current study, we report that *Pf*Sir2A possess inherent capacity to tightly bind to DNA to perform efficient deacetylation. We further uncover that *Pf*Sir2A demonstrates nucleosome-dependent deacetylase activity. it is reasoned that *Pf*Sir2A has intrinsic nucleosome binding functions, and the tight and specific interaction with nucleosome provides improved catalytic efficiency that allows *Pf*Sir2A to reduce histone acetylation.

Given its importance in controlling both the virulence and multiplicity of the parasite, *Pf*Sir2A is an attractive drug target to develop antimalarial agents. Herein, we report the discovery of endogenous and synthetic small molecule *Pf*Sir2A regulators. Free fatty acid (FFA) such as myristic acid exhibits differential regulation of the two distinct activities of *Pf*Sir2A: it activates deacetylation, but inhibits defatty-acylation. It is proposed that FFA binding-induced conformational change is responsible for the improved deacetylase efficacy. In the meantime, FFA also competes with fatty-acylated peptide substrates for the same binding site, resulting in reduced defatty-acylase activity. Furthermore, screening of a library of human sirtuin regulators leads to the discovery of two *Pf*Sir2A inhibitors: 3-TYP and nicotinamide riboside (NR). They both serve as non-competitive inhibitors to suppress *Pf*Sir2A deacetylation of synthetic peptide and physiological substrates. Taken together, our study provides not only the greatly-needed mechanistic understanding of *Pf*Sir2A regulation, but also novel chemical scaffolds for future inhibitor development.

## METHODS AND MATERIALS

### Reagents and Instruments

All reagents were purchased from Aldrich or Fisher Scientific and were of the highest purity commercially available. HPLC was performed on a Dionex Ultimate 3000 HPLC system equipped with a diode array detector using Macherey-Nagel C18 reverse-phase column. HRMS spectra were acquired with either a Waters Micromass Q-tof Ultima or a Thermo Scientific Q-Exactive hybrid Quadrupole Orbitrap.

### Synthetic Peptides

Synthetic peptides H3K9Ac: ARTKQTAR(K-Ac)STGGKAPRKQLAS, H3K9Myr: ARTKQTAR(K-Myr)STGGKAPRKQLAS, H4K16Ac: SGRGKGGKGLGKGGA(K-Ac)RHR, and p300K1024Ac: ERSTELKTEI(K-Ac)EEEDQPSTS were synthesized and purified by Genscript. The peptides were purified by HPLC to a purity >95%.

### Protein Expression and Purification

Plasmid of *Pf*Sir2A was the generous gift from Dr. Hening Lin (University of Chicago). The protein was expressed and purified according to previously published protocol.^16^ The identity of the protein was confirmed by tryptic digestion followed by LC-MS/MS analysis performed at Vermont Biomedical Research Network (VBRN) Proteomics Facility. Protein concentration was determined by Bradford assay.

### Deacetylation/Demyristoylation Assay

The *K*_m_ and *k*_cat_ of *Pf*Sir2A were measured for synthetic peptide substrates. A typical reaction was performed in 100 mM phosphate buffer pH 7.5 in a total volume of 250 µL. The reactions contained various concentrations of peptide substrate and 500 µM NAD^+^. Reactions were initiated by the addition of 5 µM (deacetylation) of 1 µM (demyristoylation) of *Pf*Sir2A and were incubated at 37 °C for 60 min (deacetylation) or 5 min (demyristoylation) before being quenched by 10% TFA. The samples were then injected on an HPLC fitted to a Macherey-Nagel Nucleosil C18 column. Acylated and deacylated peptides were resolved using a gradient of 10%–40% acetonitrile in 0.1% TFA. Chromatograms were analyzed at 215 nm. Reactions were quantified by integrating area of peaks corresponding to acylated and deacylated peptides. Rates were plotted as a function of substrate concentration and best fits of points to the Michaelis-Menten equation were performed by GraphPad Prism.

### PfSir2A Inhibition Assay

A typical reaction contained 500 µM NAD^+^, 250 µM H3K9Ac or 20 µM H3K9Myr, and varying concentrations of the small molecule inhibitor in 100 mM phosphate buffer pH 7.5. The reactions were initiated by the addition of 5 µM (deacetylation) of 1 µM (demyristoylation) of *Pf*Sir2A and were incubated at 37°C before being quenched by 10% TFA. The samples were then injected on an HPLC fitted to a Macherey-Nagel Nucleosil C18 column. Acylated and deacylated peptides were resolved using a gradient of 10%–40% acetonitrile in 0.1% TFA. Chromatograms were analyzed at 215 nm. Reactions were quantified by integrating area of peaks corresponding to acylated and deacylated peptides. Rates were plotted as a function of small molecule inhibitor concentration, and points were fitted to the following equation:

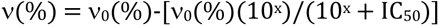

where ν(%) represents turnover rate expressed as percent enzymatic activity remaining, ν_0_(%) represents the uninhibited turnover rate expressed as an enzymatic activity of 100%. The variable x represents the log[inhibitor] in nanomolar. IC_50_ values were derived from this equation.

### Mode of Inhibition Analysis

Peptide titration reactions containing 0, 10, 100, or 1000 µM inhibitor were incubated with 500 µM NAD^+^, varying concentrations of H3K9Ac or H3K9Myr in 100 mM phosphate buffer pH 7.5. NAD^+^ titration reactions containing 0, 10, 100, or 1000 µM inhibitor were incubated with 250 µM H3K9Ac or 20 µM H3K9Myr, varying concentrations of NAD^+^ in 100 mM phosphate buffer pH 7.5. Reactions were initiated by the addition of 5 µM (deacetylation) of 1 µM (demyristoylation) of *Pf*Sir2A and were incubated at 37 °C before being quenched by 10% TFA. The samples were then injected on an HPLC fitted to a Macherey-Nagel Nucleosil C18 column. Acylated and deacylated peptides were resolved using a gradient of 10%– 40% acetonitrile in 0.1% TFA. Chromatograms were analyzed at 215 nm. Reactions were quantified by integrating area of peaks corresponding to acylated and deacylated peptides. From the Michaelis-Menten plots, *K*_m_ and *k*_cat_ were calculated for different concentrations of the inhibitor.

### Electrophoretic Mobility Shift Assay (EMSA)

dsDNA was incubated with varying concentrations of *Pf*Sir2A (0 to 20 µM) in DNA binding buffer (20 mM Tris-HCl, pH 8.0, 150 mM NaCl, 2 mM DTT, 5% glycerol). The samples were incubated on ice for 20 min. The samples were then resolved on a native 4∼20% TBE gel at 4 °C. After SYBR safe staining, the visualization and quantification of the gels were carried out using a Biorad ChemiDoc MP imaging system and Image Lab software. Any species migrating at a slower electrophoretic rate than free DNA were considered as a DNA-protein complex. Relative complex formation was plotted as a function of protein concentration and fitted to the equation: *f* = *f*_max_ [*Pf*Sir2A]^h^/(*K*_d_ + [*Pf*Sir2A]^h^), where *f* is the fractional saturation, *f*_max_ is maximum complex formation, [*Pf*Sir2A] is the protein concentration, h is the Hill slope, and *K*_d_ reflects the binding affinity of the proteins to the DNA construct.

### Microscale Thermophoresis (MST) Binding Assay

The MST experiments were performed using a Monolith NT.115 instrument (NanoTemper Technologies). The purified recombinant His-tagged *Pf*Sir2A was labeled by the RED-tris-NTA 2^nd^ generation dye (NanoTemper Technologies). The protein concentration was adjusted to 200 nM in PBS-T buffer, while the dye concentration was set to 100 nM. Equal volumes (100 µL) of protein and dye solutions were mixed and incubated at room temperature in the dark for 30 min. Sixteen solutions of ssDNA with decreasing concentrations were prepared in the MST buffer by serial half-log dilution. Each solution was mixed with an equal volume of the labeled *Pf*Sir2A. The binding affinity analysis was performed using standard MST capillaries, 40% excitation power (Nano-RED) and medium MST power at room temperature. The *K*_d_ value was determined with the MO.Affinity Analysis software (NanoTemper Technologies), using three independent MST measurements.

### Cell Culture

HEK293 cells were cultured in DMEM supplemented with 10% fetal bovine serum (FBS), 100 U/mL penicillin and 100 mg/mL streptomycin. Cells were maintained in a humidified 37°C incubator with 5% CO_2_.

### Metabolic Labeling

Cells were treated with either 20 µM azido myristic acid or DMSO (vehicle control) for 18 h. Cells were harvested, re-suspended in PBS, and pelleted at 1,000 g for 5 min at 4 °C. The cell pellet was then dissolved in RIPA buffer (Thermo Scientific) with Halt protease phosphatase inhibitor cocktail (Thermo Scientific). The lysate was centrifuged at 15,000 g for 5 min at 4 °C. The supernatant was used for the defatty-acylation analysis.

### Defatty-acylation Assay

Cell lysate was incubated with 500 µM NAD^+^, 10 µM *Pf*Sir2A with or without calf thymus DNA at 37 °C for 1 h. The samples were then incubated with 15 mM iodoacetamide at room temperature for 30 min before TAMRA-DBCO was added. The samples were incubated at room temperature for another 30 min. Subsequently, the samples were heated with NH_2_OH (60 mM, pH 7.2) at 95 °C for 7 min before being resolved by SDS-PAGE. The destained gel was analyzed with in-gel fluorescence scanning using a Biorad ChemiDoc MP imager (excitation at 532 nm, 580 nm cut-off filter and 30 nm band-pass).

### Western Blot

The samples were resolved on a 10% SDS-PAGE gel and transferred to Immobilon PVDF transfer membrane (Biorad). The blot was blocked with 5% nonfat milk, probed with primary antibody, washed with TBST, followed by incubation with anti-rabbit HRP-conjugated secondary antibody. The signal was then detected by Clarity western ECL substrate (Biorad).

## RESULTS AND DISCUSSION

### Characterization of PfSir2A activity in vitro

We have expressed and purified *Pf*Sir2A with an *N*-terminal His-tag. It can deacetylate a panel of peptide substrates.^14^ The kinetic constants for *Pf*Sir2A-mediated deacetylation under steady-state conditions at pH 7.5 have been determined (**Table 1**). The recombinant enzyme is also able to deacetylate H3K9Ac in calf thymus histones (**Figures 1A** and **1B**). In *in vitro* setting, *Pf*Sir2A demonstrates weak deacetylase activity towards the synthetic peptide substrates, consistent with a previous report.^16^ In contrast, the catalytic efficiency (*k*_cat_/*K*_m_) of demyristoylation is 4400-fold higher than that of the deacetylation (5140 M^-1^s^-1^ *vs.*1.15 M^-1^s^-1^, **Table 1**), confirming the robust defatty-acylase activity of *Pf*Sir2A.^16^

**Figure 1.**
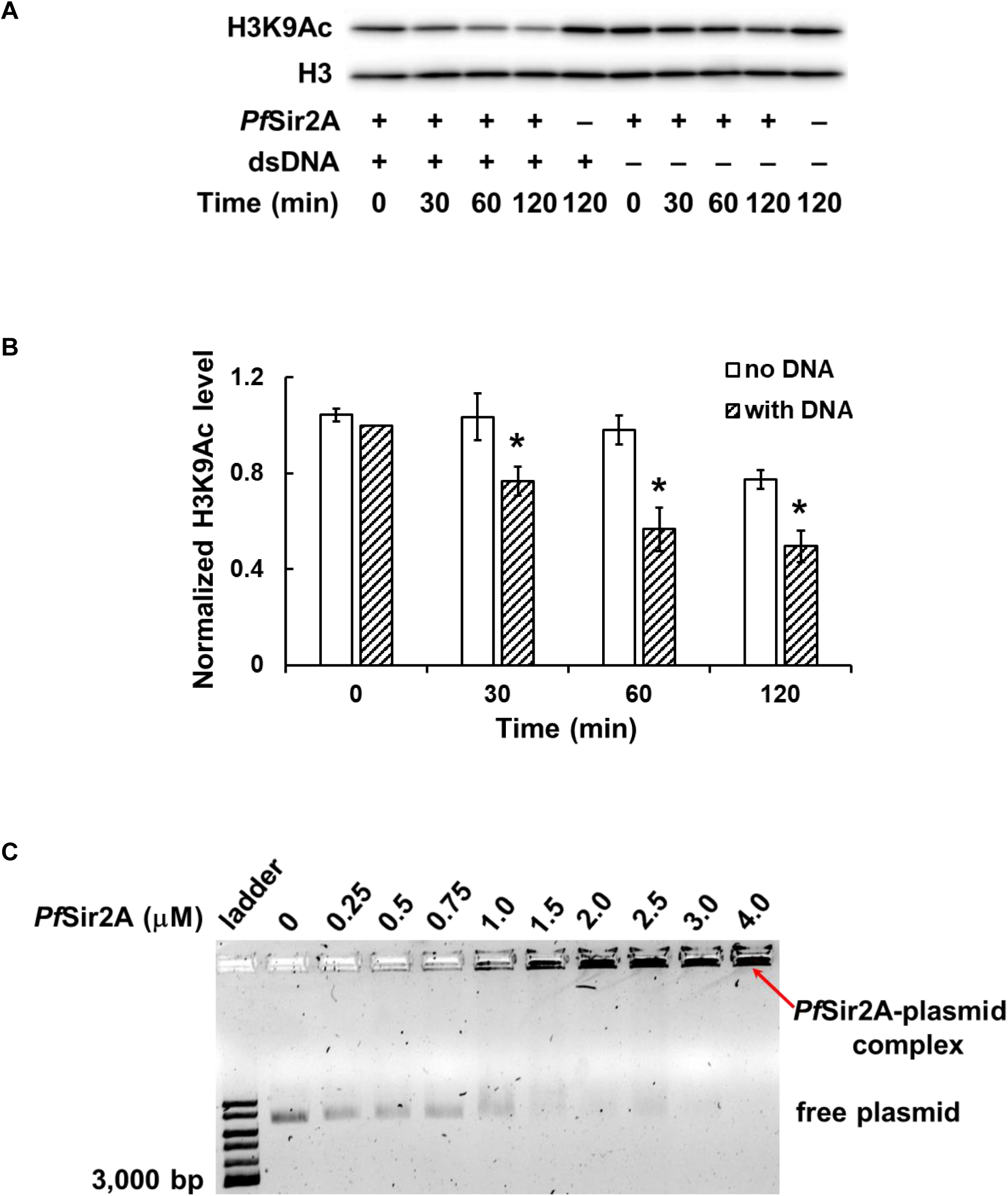
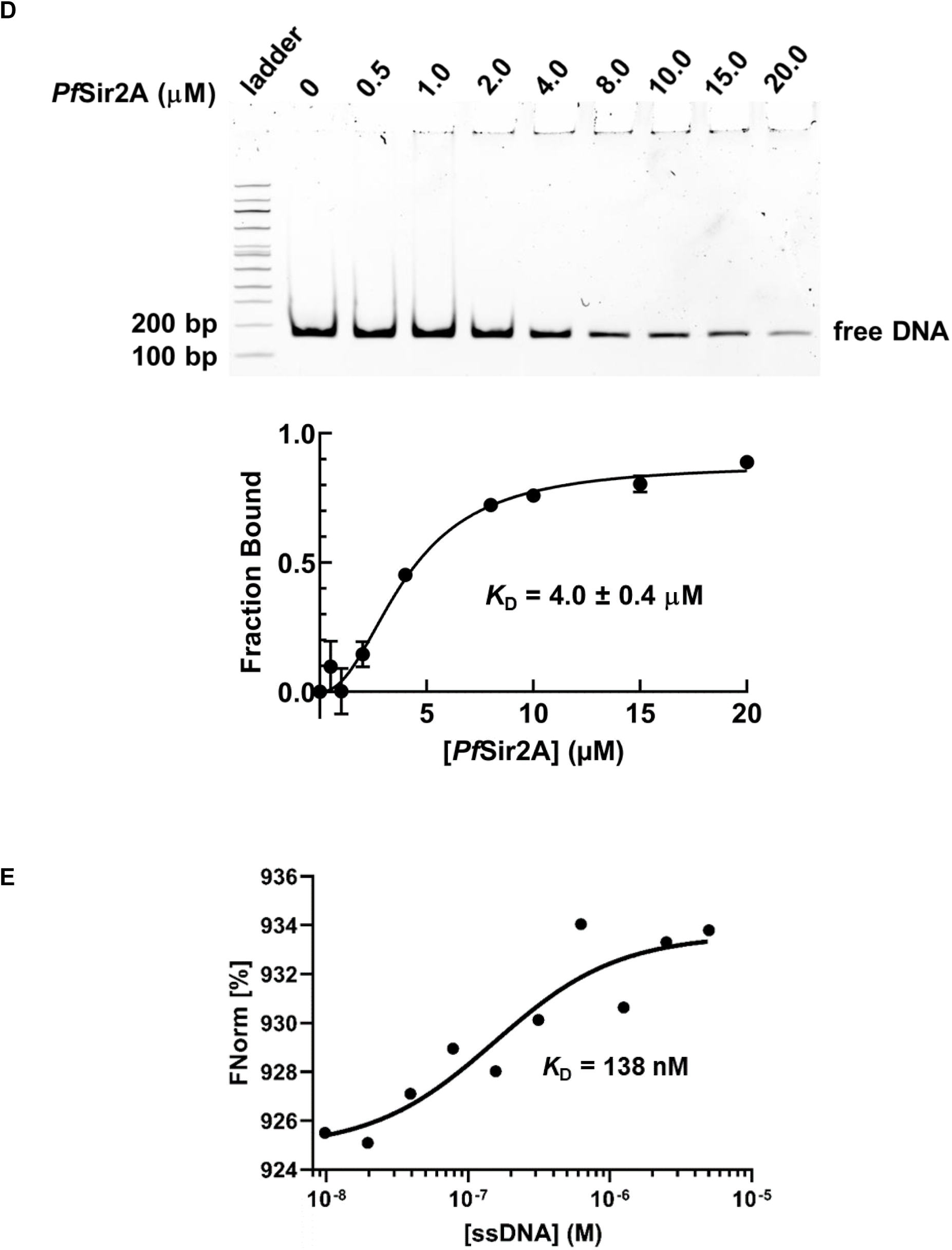
Analysis of *Pf*Sir2A interaction with DNAs. (A) Representative western blots showing the deacetylase activity of *Pf*Sir2A on calf thymus histone in the presence or absence of dsDNA; (B) Quantification of the western blot results. Statistical significance was determined by a Student’s *t*-test: * *p* < 0.05 *vs* no dsDNA control; (C) Representative gel image of the EMSA analysis of the interaction of *Pf*Sir2A with a circular plasmid; (D) EMSA analysis of the interaction of *Pf*Sir2A with the nucleosome assembly 601 sequence DNA. Top: representative gel image; Bottom: binding affinity analysis; (E) Determination of binding affinity of *Pf*Sir2A to ssDNA using MST. The sequence of the ssDNA is GAAGTCGGGGATGGCAGAGGCAGTGCT.

**Table 1.** Recombinant *Pf*Sir2A steady state parameters.

| Substrate<sup>a</sup> | Additive | $K_m$ (μM) | $k_{cat}$ ($10^{-4}s^{-1}$) | M.A.<sup>f</sup> |
| --- | --- | --- | --- | --- |
| H3K9Ac<sup>b</sup> | | 556 ± 54 | 6.4 ± 0.8 | 1.0 |
| H3K9Ac | dsDNA<sup>e</sup> | 175 ± 22 | 8.2 ± 0.5 | 4.1 |
| H4K16Ac<sup>c</sup> | | 195 ± 42 | 4.3 ± 0.5 | |
| p300K1040Ac<sup>d</sup> | | 102 ± 21 | 11.4 ± 1.2 | |
| H3K9Myr<sup>b</sup> | | 4.9 ± 1.1 | 252 ± 17 | 1.0 |
| H3K9Myr | dsDNA | 0.95 ± 0.3 | 305 ± 28 | 6.2 |

<sup>a</sup> At saturating NAD<sup>+</sup> concentration; <sup>b</sup> Sequence: H3(1-22); <sup>c</sup> Sequence: H4(1-19);
<sup>d</sup> Sequence: p300(1034-1053); <sup>e</sup> [dsDNA] = 200 ng/ μL;
<sup>f</sup> M.A.: Maximal Activation = $k_{cat}(act.) * K_m(unact.) / (k_{cat}(unact.) * K_m(act.))$ .

### PfSir2A activities are activated by DNA

Inspired by the early suggestions that *Pf*Sir2A is a telomeric binding factor,^7, 10, 17^ we performed a simple *Pf*Sir2A-DNA binding experiment. The ability of *Pf*Sir2A to bind to DNA was evaluated using the electrophoretic mobility shift assay (EMSA) with a circular plasmid (**Figure 1C**): in the samples containing *Pf*Sir2A and plasmid (lanes 3 to 11 ), the formation of a *Pf*Sir2A-DNA complex can be readily detected (indicated by the red arrow). Similarly, *Pf*Sir2A demonstrates tight binding to the nucleosome assembly 601 sequence DNA^18^ (a 147 bp dsDNA) with a *K*_D_ value of 4.0 ± 0.4 µM (**Figure 1D**). *Pf*Sir2A also binds to ssDNA (*K*_d_ = 138 nM) as determined by microscale thermophoresis (MST) assay (**Figure 1E**). The interaction between *Pf*Sir2A and nucleic acids is sequence-independent.

These initial findings prompted us to investigate whether the deacetylase activity of *Pf*Sir2A can be activated upon binding to DNA. Indeed, in the presence of calf thymus DNA (dsDNA sheared to an average size less than 2000 bp), the catalytic efficiency of deacetylation exhibits a 4-fold improvement towards the peptide substrate (**Table 1**), owing to a reduced *K*_m_ and an increased *k*_cat_. Furthermore, the effect of DNA on physiological *Pf*Sir2A substrate was also assessed. Calf thymus histone was incubated with recombinant *Pf*Sir2A and NAD^+^ with or without dsDNA. The acetylation level of H3K9 was then probed using western blot. The addition of dsDNA caused a significant reduction of H3K9Ac compared to the no DNA controls (**Figures 1A** and **1B**). To our knowledge, these results represent the first direct evidence that the enzymatic activity of *Pf*Sir2A can be stimulated upon binding to DNA strands.

Studies from the last two decades have suggested that *Pf*Sir2A is a promiscuous enzyme with multiple enzymatic activities. In addition to deacetylation^14^ and mono-ADP-ribosylation,^13^ *Pf*Sir2A also harbors defatty-acylation activity.^16^ As *Pf*Sir2A is poised at the intersection between chromatin modulation and genome maintenance, understanding how this novel activity is regulated is highly relevant. In our preliminary study, a myristoylated synthetic H3K9 peptide (H3K9Myr) was used as the substrate for the characterization of the defatty-acylase activity of *Pf*Sir2A. The inclusion of dsDNA caused a marked reduction of *K*_m_ (from 4.9 to 0.95 µM, **Table 1**), leading to a 6-fold activation of the demyristoylase activity.

To examine if lysine fatty-acylation of cellular proteins can be modulated by *Pf*Sir2A, azido myristic acid (Az-myr) was used for the metabolic labeling of cells. As shown in **Figure 2A**, HEK293 cells were treated with Az-myr (20 µM) which can be metabolically incorporated into fatty-acylated, especially *N-* and *S*-myristoylated, cellular proteins. The cell lysate was then incubated with *Pf*Sir2A in the presence or absence of dsDNA. Subsequently, the labeled proteins were covalently tethered to TAMRA-DBCO *via* strain-promoted “click” conjugation. NH_2_OH was used to remove the potential cysteine fatty-acylation. Thus, the in-gel fluorescence signal will mainly represent the lysine fatty-acylation level. Compared to the control cell lysate, Az-myr-treated cell lysate demonstrated significantly increased labeling intensity (**Figure 2B**). Treatment with *Pf*Sir2A in the presence of NAD^+^ reduced the fluorescence signals of numerous proteins, confirming the previous hypothesis that *Pf*Sir2A can remove lysine fatty-acylation from endogenous protein targets.^16^ The addition of dsDNA to the sample caused further signal reduction of several highlighted proteins. Although the chemical nature of lysine acylation has been known to regulate the level of *Pf*Sir2A catalytic efficiency,^16^ these substrate preferences have never been studied in combination with additional modulators such as nucleic acids. Our data suggest that DNA-complexed *Pf*Sir2A displays enhanced defatty-acylase activity.

**Figure 2.**
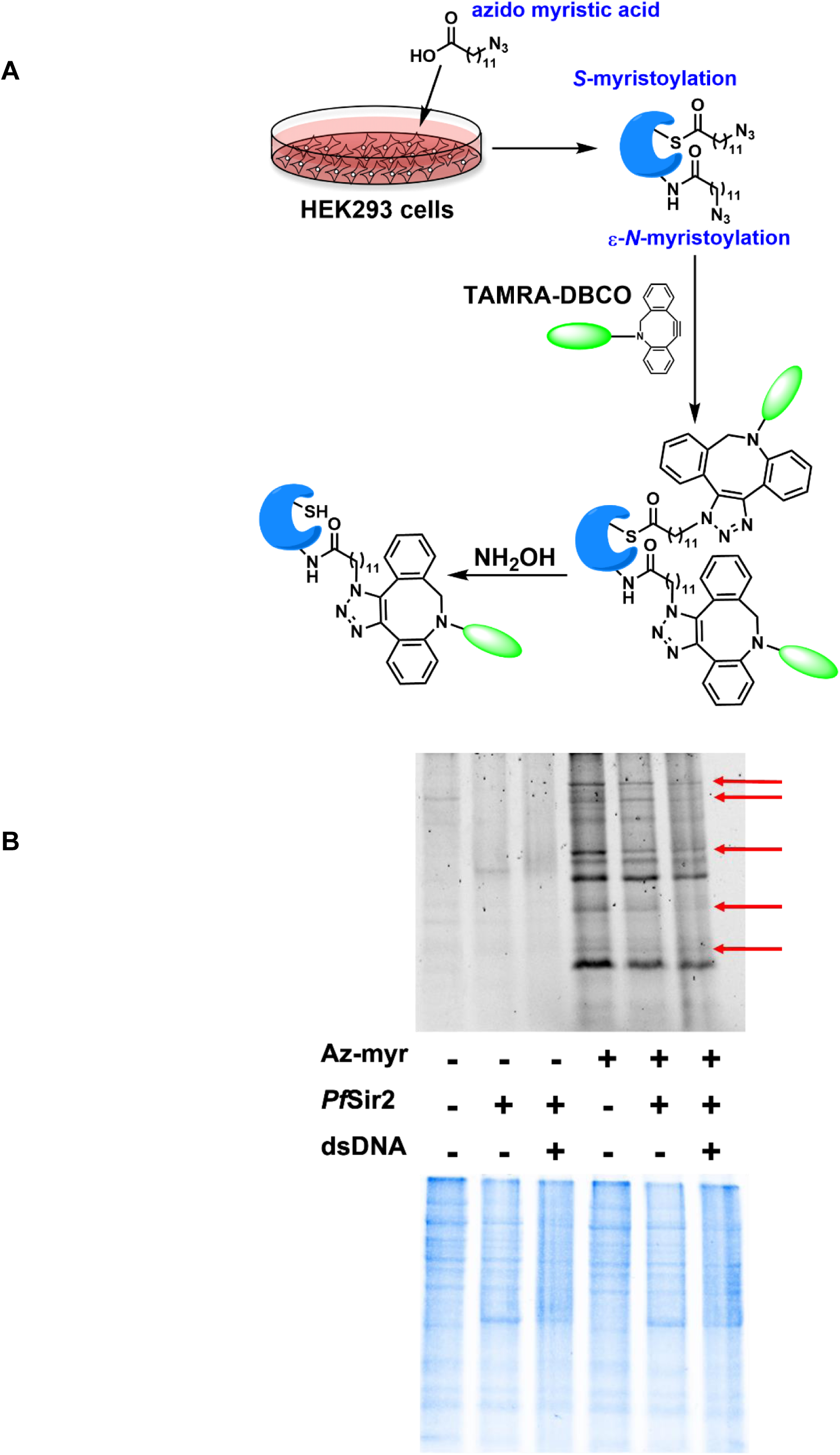
Defatty-acylase activity of *Pf*Sir2A can be activated by dsDNA. (A) Schematic representation of the metabolic labeling; (B) Defatty-acylation of cellular proteins by *Pf*Sir2A. The lysate from Az-myr treated cells showed increased labeling compared to the control cells. Incubation of the lysate with *Pf*Sir2A in the presence of dsDNA reduced the labeling intensity of several proteins compared to the no DNA control. Arrows highlight proteins with decreased fluorescence due to dsDNA exposure.

### Differential regulation of PfSir2A activity by free fatty acid

A co-crystal structure of *Pf*Sir2A with NAD^+^ and a bound H3H9Myr peptide provides the structural basis of the preferred defatty-acylase activity.^16^ *Pf*Sir2A has a long hydrophobic tunnel to accommodate the long chain fatty-acyl groups (**Figure 3A**). Structural alignment of *Pf*Sir2A-AMP (pdb: 3JWP) and *Pf*Sir2A-NAD^+^-H3K9Myr (pdb: 3U3D) suggests that binding of the H3K9Myr peptide drives the zinc-binding domain to rotate clockwise to the Rossman fold domain, resulting in *Pf*Sir2A moving from an inactive open state to a productive closed state.^19^ It is reasonable to speculate that the hydrophobic tunnel of *Pf*Sir2A allows it to bind to free fatty acids (FFAs) for conformational rearrangement and the stimulation of its deacetylase activity.

**Figure 3.**
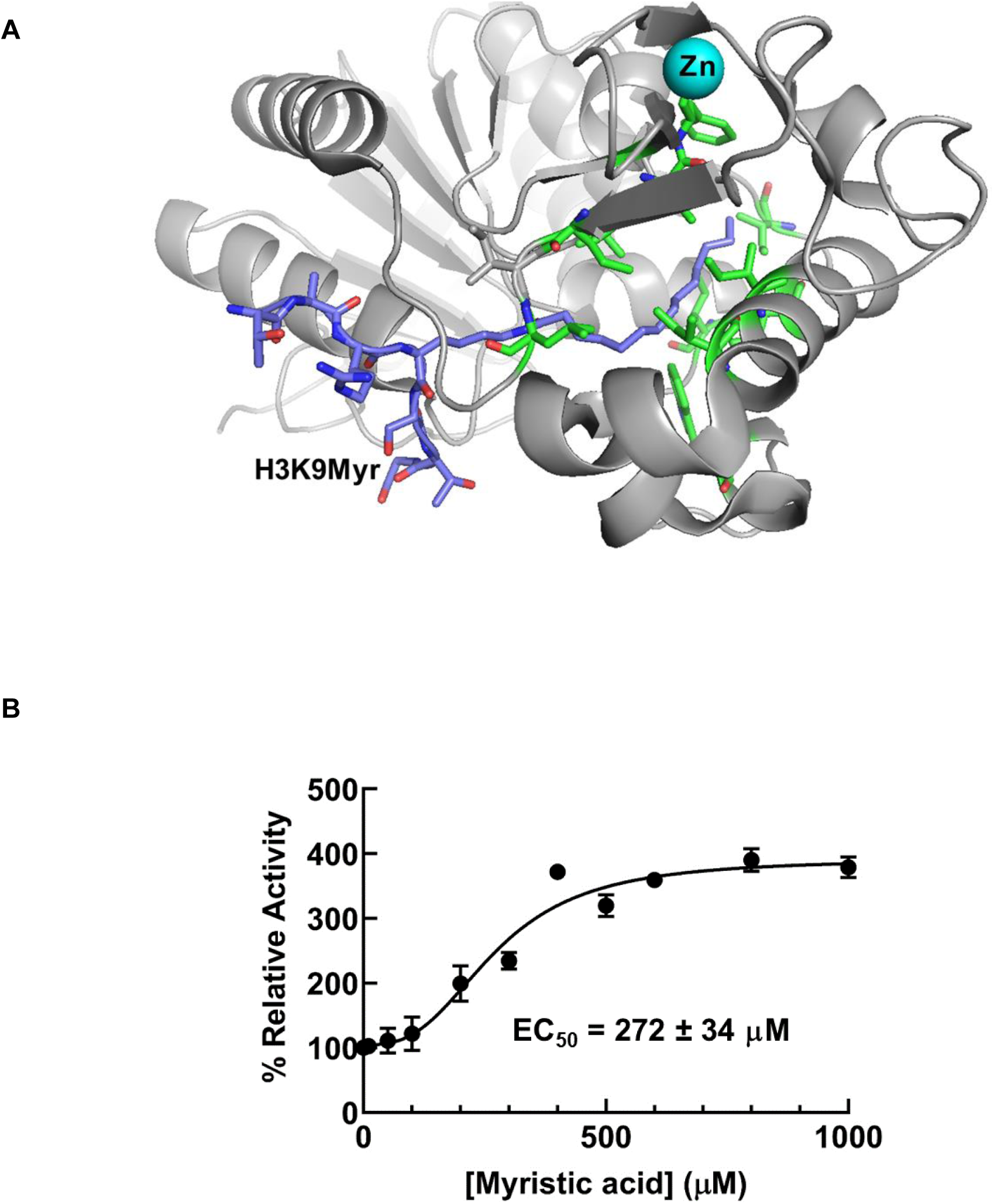
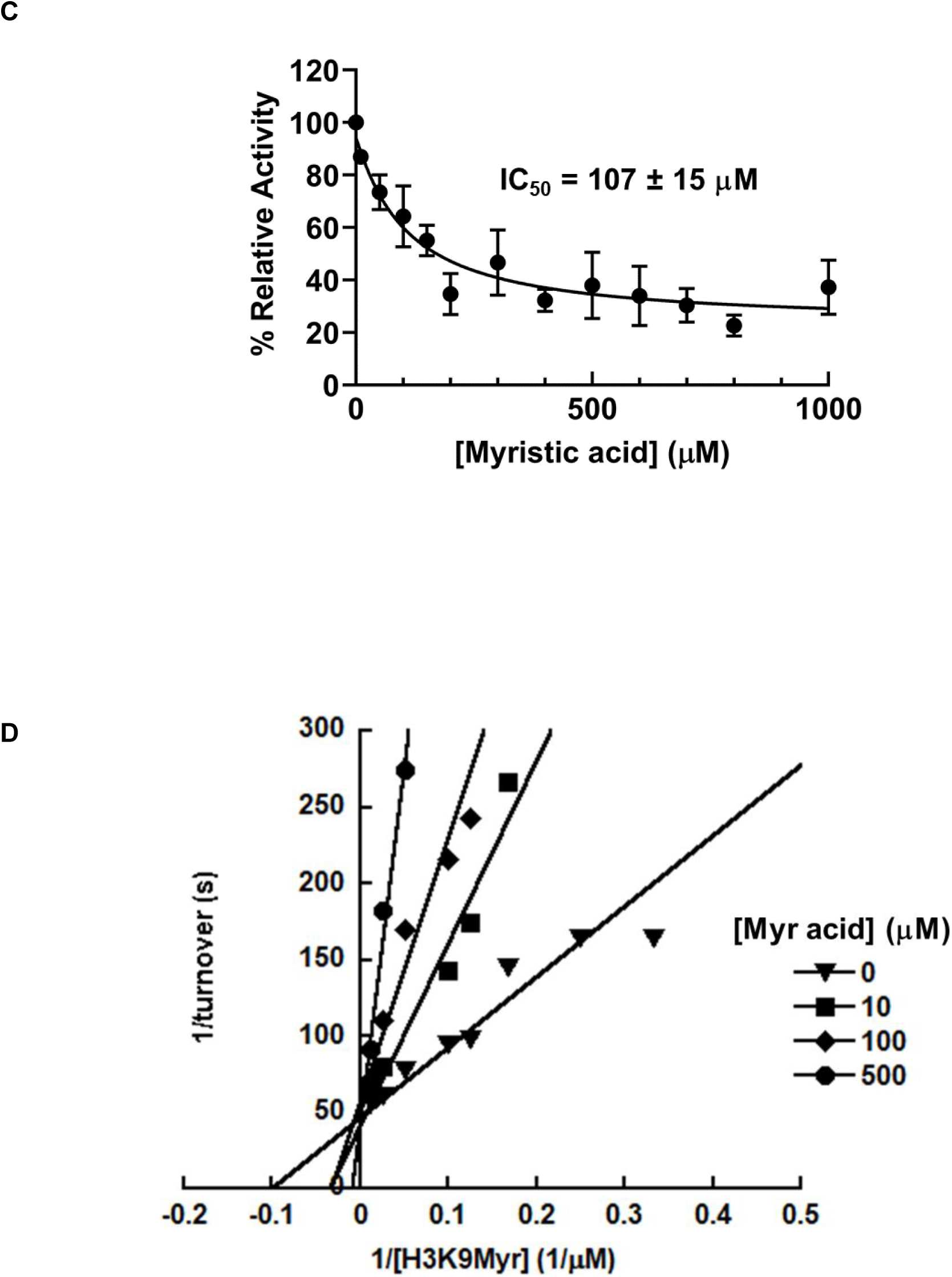
Differential regulation of *Pf*Sir2A activity by myristic acid. (A) *Pf*Sir2A in complex with H3K9Myr (pdb: 3U3D). H3K9Myr: purple; zinc: cyan; hydrophobic amino acids in the binding tunnel: green; (B) Activation of *Pf*Sir2A deacetylase activity by myristic acid; (C) Inhibition of *Pf*Sir2A demyristoylase activity by myristic acid; (D) Double-reciprocal plot showing that myristic acid is a competitive inhibitor of *Pf*Sir2A’s demyristoylase activity.

To assess if FFAs can activate *Pf*Sir2A deacetylation, the enzyme was incubated with NAD^+^ and H3K9Ac peptide in the presence of myristic acid. The concentrations of myristic acid were maintained below its critical micelle concentration (CMC).^20^ Indeed, the deacetylation was stimulated by myristic acid in a concentration-dependent fashion (**Figure 3B**): a maximum of 4-fold activation with an EC_50_ of 272 ± 34 µM. The EC_50_ value is well within the physiological concentrations of FFAs in humans.^21^

The ability of myristic acid to activate *Pf*Sir2A deacetylation supports our initial hypothesis that myristic acid occupies the same binding pocket as the myristoylated peptide substrate. To further confirm this, the effect of myristic acid on the demyristoylase activity of *Pf*Sir2A was also examined. As shown in **Figure 3C**, demyristoylation of H3K9Myr was inhibited by myristic acid in a dose-dependent manner with an IC_50_ value of 107 ± 15 µM. The mode of inhibition was then investigated by steady-state inhibition analysis. Double reciprocal plot showed that myristic acid was competitive with H3K9Myr peptide as evidenced by a series of lines that intersect at the y-axis (**Figure 3D**), consistent with the notion that FFA competes with myristoylated peptide for the same binding site.

*var* gene transcription is epigenetically regulated by *Pf*Sir2A,^3,8^ a NAD^+^-dependent histone deacetylase. Given recent evidence supporting that FFAs can reduce histone acetylation levels at specific loci,^22, 23^ we hypothesize that metabolism and epigenetic regulation of gene transcription may be controlled by FFA availability. The parasites acquire FFA through either *de novo* synthesis or scavenging from the host cells.^24^ Some recent studies reveal that *P. falciparum* senses nutrient availability to induce epigenetic reprogramming.^25, 26^ When FFAs are abundant, the deacetylase activity of *Pf*Sir2A can be stimulated for targeted histone deacetylation and *var* gene silencing. FFAs may also serve as an intrinsic regulatory factor allowing *Pf*Sir2A to switch from a defatty-acylase to a deacetylase, although the biological significance of *Pf*Sir2A-catalyzed defatty-acylation remains elusive.

### PfSir2A demonstrates nucleosome-dependent activity

Despite a plethora of reported *in vivo* functions, the biochemical aspects of *Pf*Sir2A activity and regulation remain poorly understood. The recombinant *Pf*Sir2A is found to be rather inactive, but activatable toward acetylated substrates by complexation with DNAs. This suggests the existence of a more potent, but previously uncharacterized, *Pf*Sir2A activation mechanism. Considering the chromatin association and function,^7, 8^ it is reasonable to hypothesize nucleosomes as potential *Pf*Sir2A activators.

To study *Pf*Sir2A-nucleosome interactions, a highly purified recombinant human 601-positioned nucleosome (EpiCypher) was used. In this nucleosome, histone H3 contains an acetyllysine at position 9 (H3K9Ac dNuc). The binding interaction was assessed using the EMSA. The titration of recombinant *Pf*Sir2A into nucleosome resulted in the formation of a higher molecular weight band (**Figure 4A**). H3K9Ac dNuc effectively interacts with *Pf*Sir2A with a *K*_D_ value of 1.4 ± 0.2 µM (**Figure 4B**). To further characterize if binding to nucleosome promotes more efficient histone deacetylation, the ability of *Pf*Sir2A to deacetylate H3K9Ac dNuc was evaluated using western blot. The enzyme showed enhanced deacetylation of nucleosome substrate as compared to free histone proteins (**Figures 4C** and **4D)**. Our data suggest that *Pf*Sir2A harbors the intrinsic capacity to bind tightly and remove acetyl groups efficiently from nucleosomes. Currently, we are using a unique combination of HPLC-based enzyme activity assays, EMSA, as well as site-directed mutagenesis and truncation analysis to investigate the molecular features required for the intimate *Pf*Sir2A-nucleosome interactions, and how these interactions contribute to the efficient removal of acetyl marks from histone lysines.

**Figure 4.**
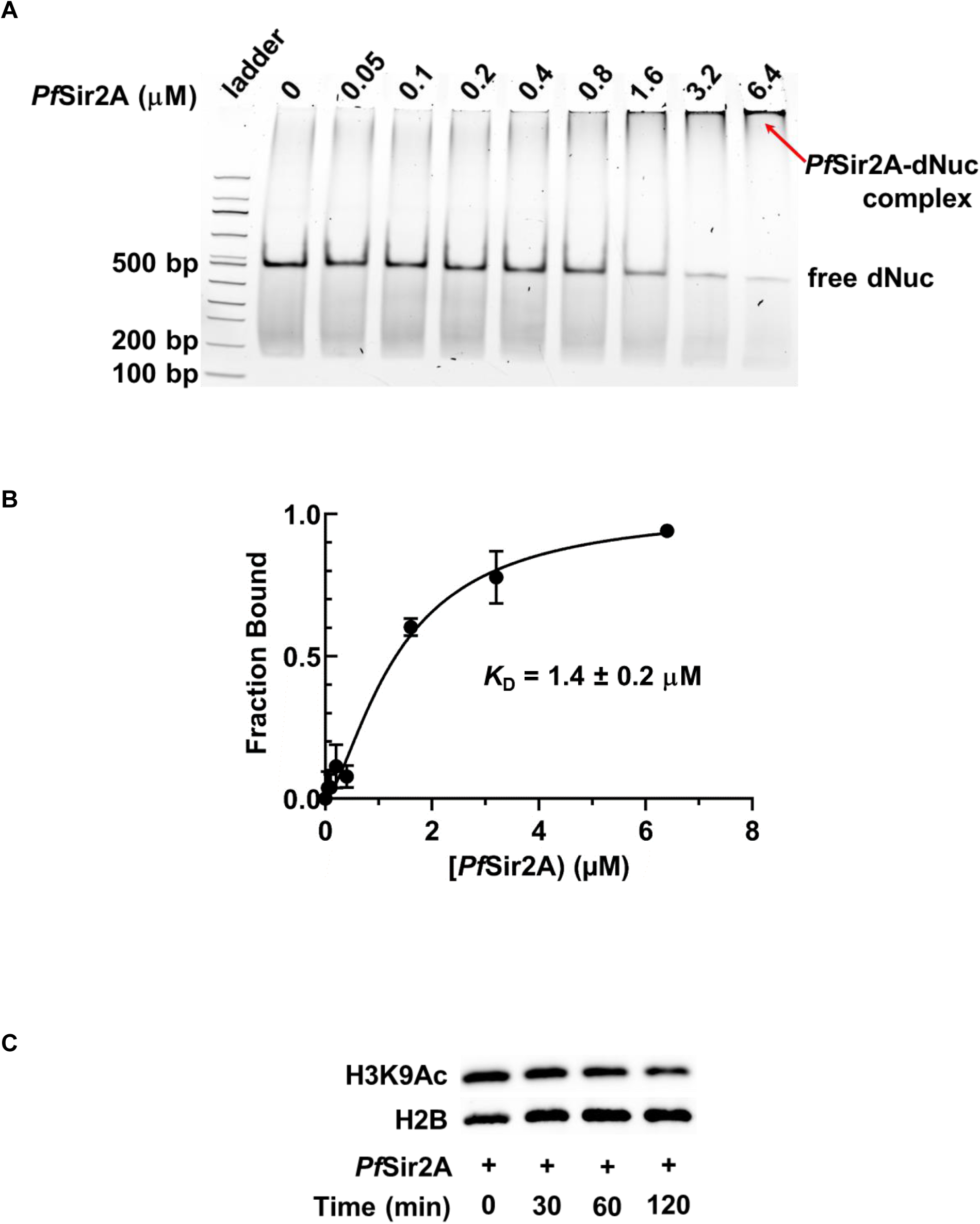
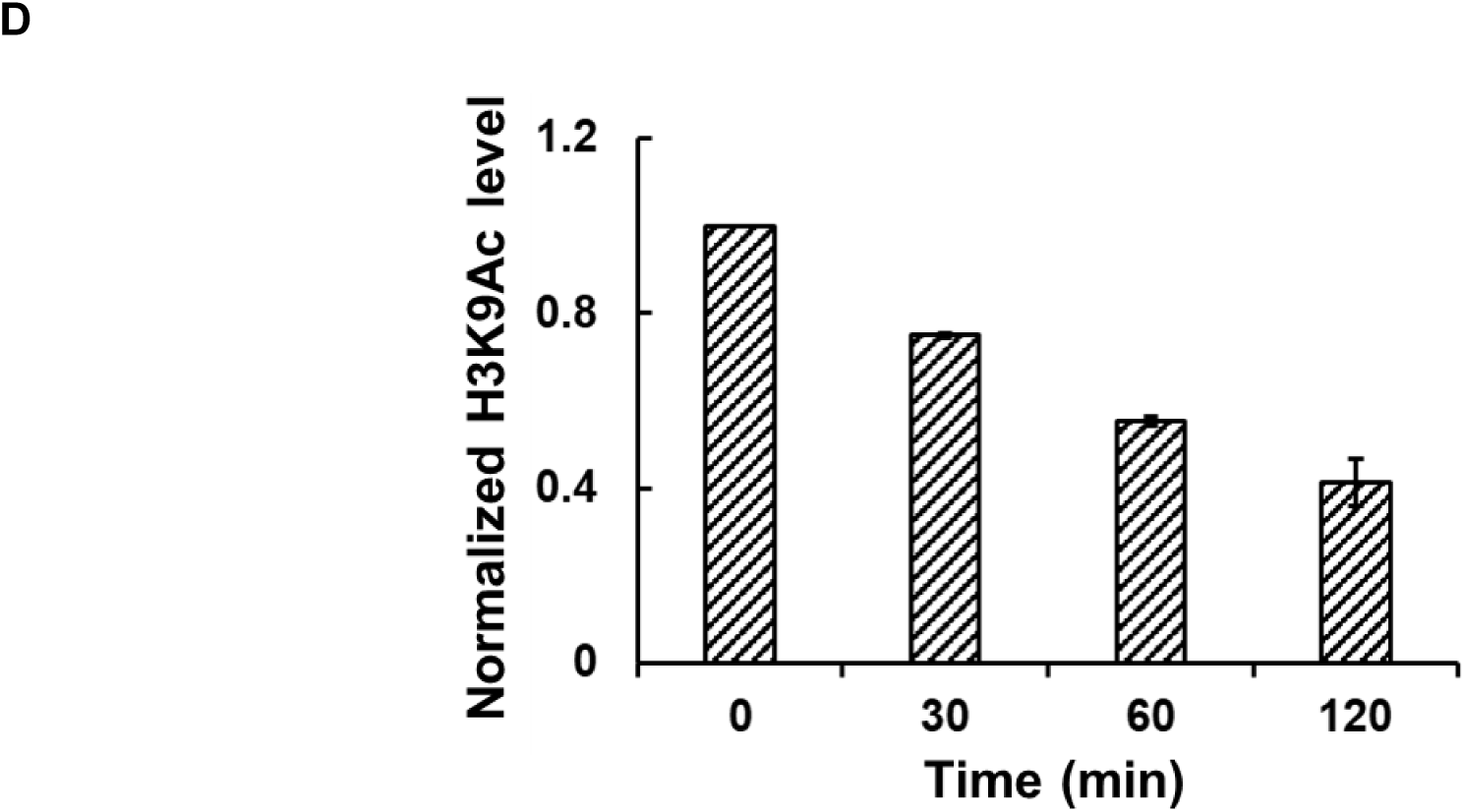
*Pf*Sir2A demonstrates nucleosome-dependent deacetylase activity. (A) EMSA analysis of *Pf*Sir2A interactions with H3K9Ac dNuc; (B) Binding affinity analysis. The *K*_D_ was determined to be 1.4 ± 0.2 µM; (C) Western blot analysis of *Pf*Sir2A deacetylation of H3k9Ac dNuc; (D) Quantification of the western blot results in (C). The H3K9Ac level was normalized by the amount of histone H2B protein.

### Repurposing human sirtuin regulators as PfSir2A inhibitors

Although combination therapies have succeeded in reducing the global burden of malaria, multidrug resistance is emerging worldwide. Innovative antimalarial drugs that kill all-life-cycle stages of parasites are urgently needed. *Pf*Sir2A controls both the virulence and multiplicity of the parasite, and disruption of this gene results in loss of mono-allelic expression of *var* genes. Thus, this epigenetic regulator is an attractive drug target for overcoming multidrug resistance.

Currently, there are no *Pf*Sir2A-selective inhibitors. Nicotinamide (NAM) is the physiological Sir2 inhibitor.^27,28^ A recent study indicates that NAM inhibits *Pf*Sir2A deacetylase activity *in vivo*.^29^ However, this compound can be directly incorporated into metabolic pathways and shows broad-spectrum inhibition against Sir2s. A lysine-based tripeptide analog has been shown to inhibit recombinant *Pf*Sir2A activity, and inhibit the intra-erythrocytic parasite growth.^30^ However, this compound exhibits comparable potency against human SIRT1. An ideal *Pf*Sir2A inhibitor should have a novel scaffold that allows it to target *Pf*Sir2A in a selective manner.

The development of human sirtuin regulators has grown exponentially during the last two decades, with several compounds in clinical trials for various diseases.^31^ These prior studies not only provide chemical entities for drug discovery, but also insight into sirtuin structure, selectivity, and biological functions. All of these can be harnessed for the development of *Pf*Sir2A inhibitors. To this end, we screened a commercially available library of 28 human sirtuin regulators (Selleckchem) at 10, 100, and 1000 µM against recombinant *Pf*Sir2A. Using the HPLC-based assay described in “Methods and Materials”, the effect of these compounds of *Pf*Sir2A-catalyzed deacetylation was evaluated. 3-TYP and nicotinamide riboside (NR) demonstrated concentration-dependent inhibition of *Pf*Sir2A deacetylase activity with more than 50% inhibition at 1 mM concentration (**Figure 5A**). The IC_50_ value of NR, a water soluble compound, was determined to be 155.2 ± 14.9 µM (**Figure 5B**). Due to its limited solubility, the IC_50_ of 3-TYP was estimated to be 59.6 ± 3.4 µM (**Figure 5C**).

**Figure 5.**
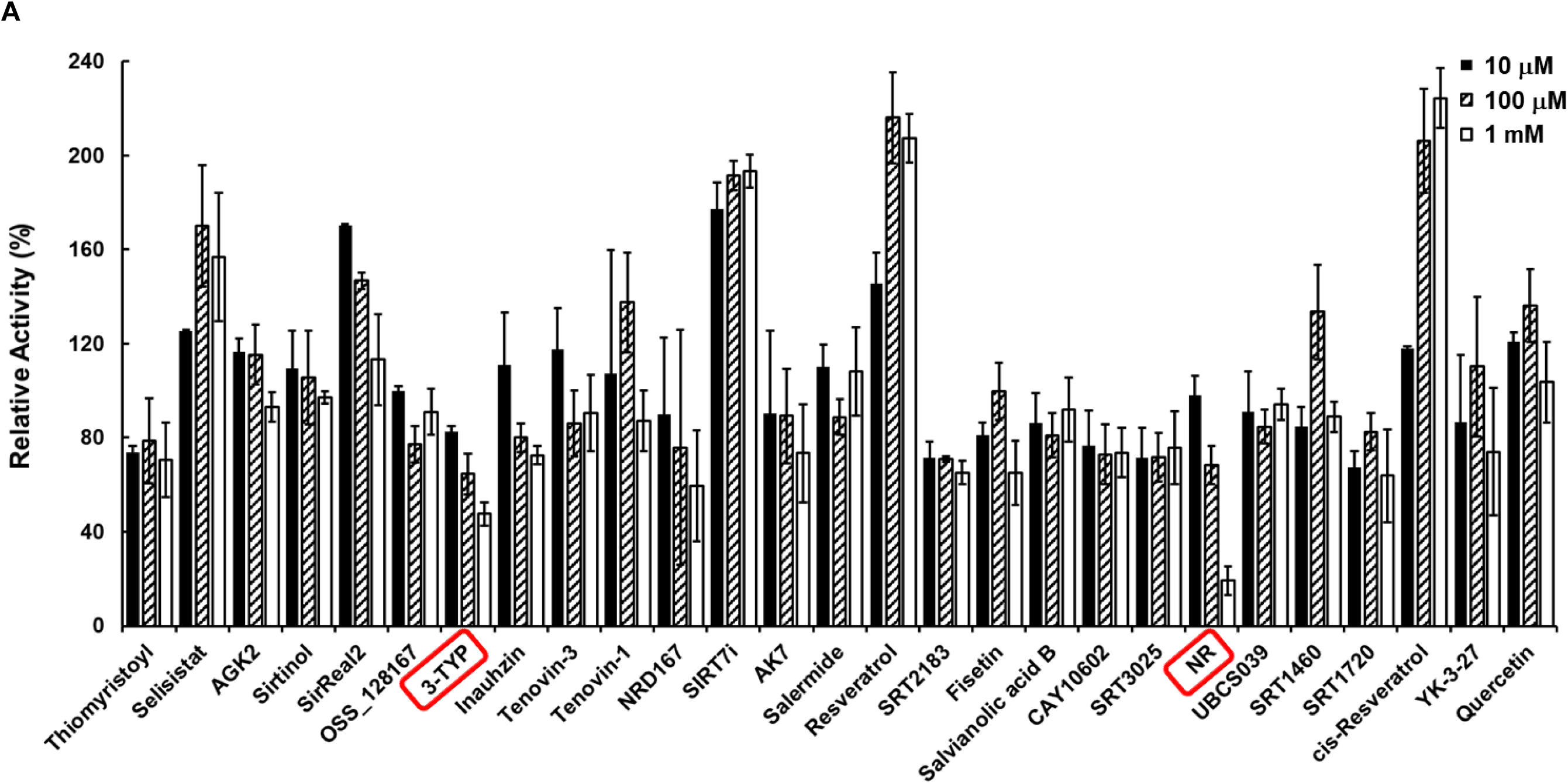
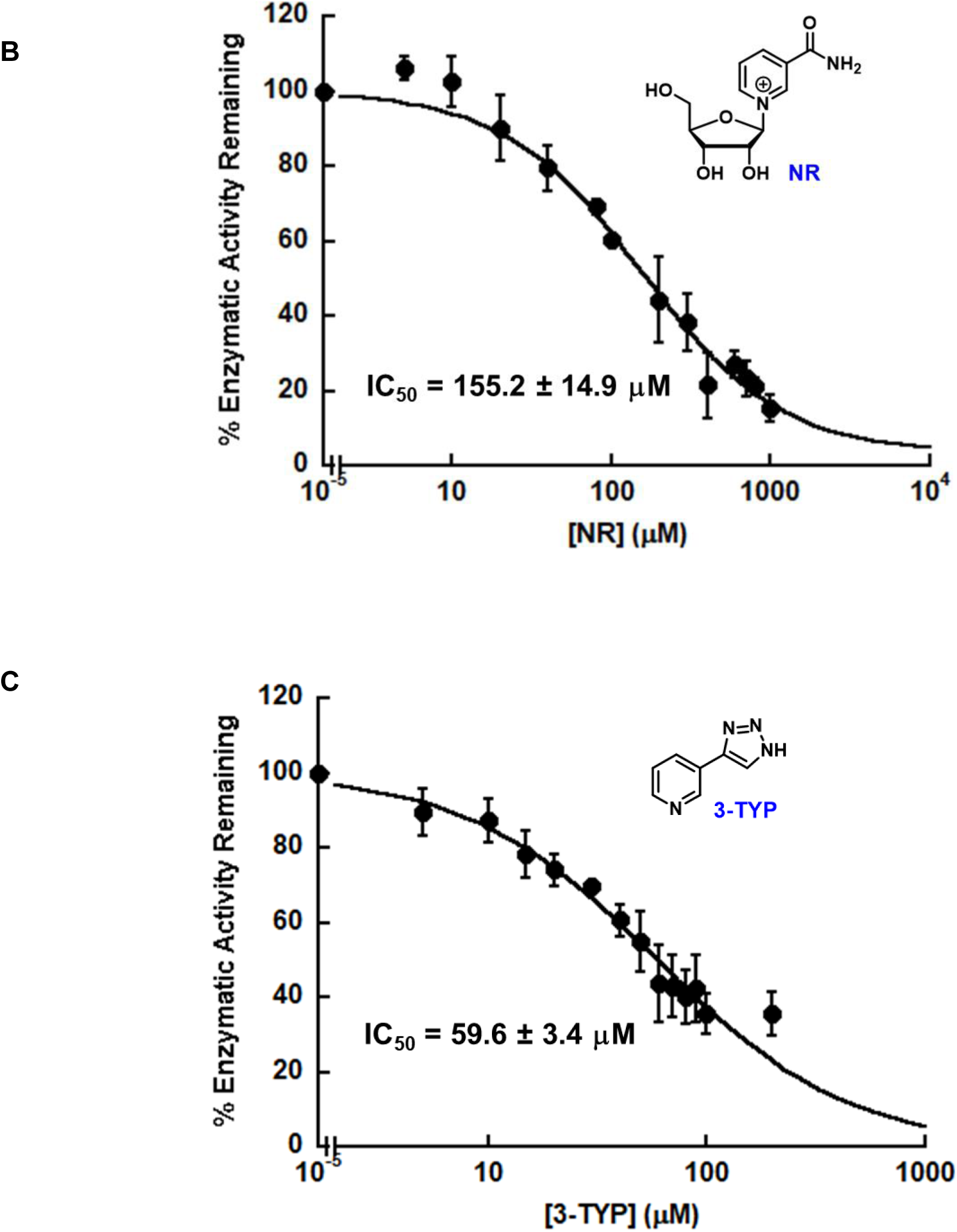
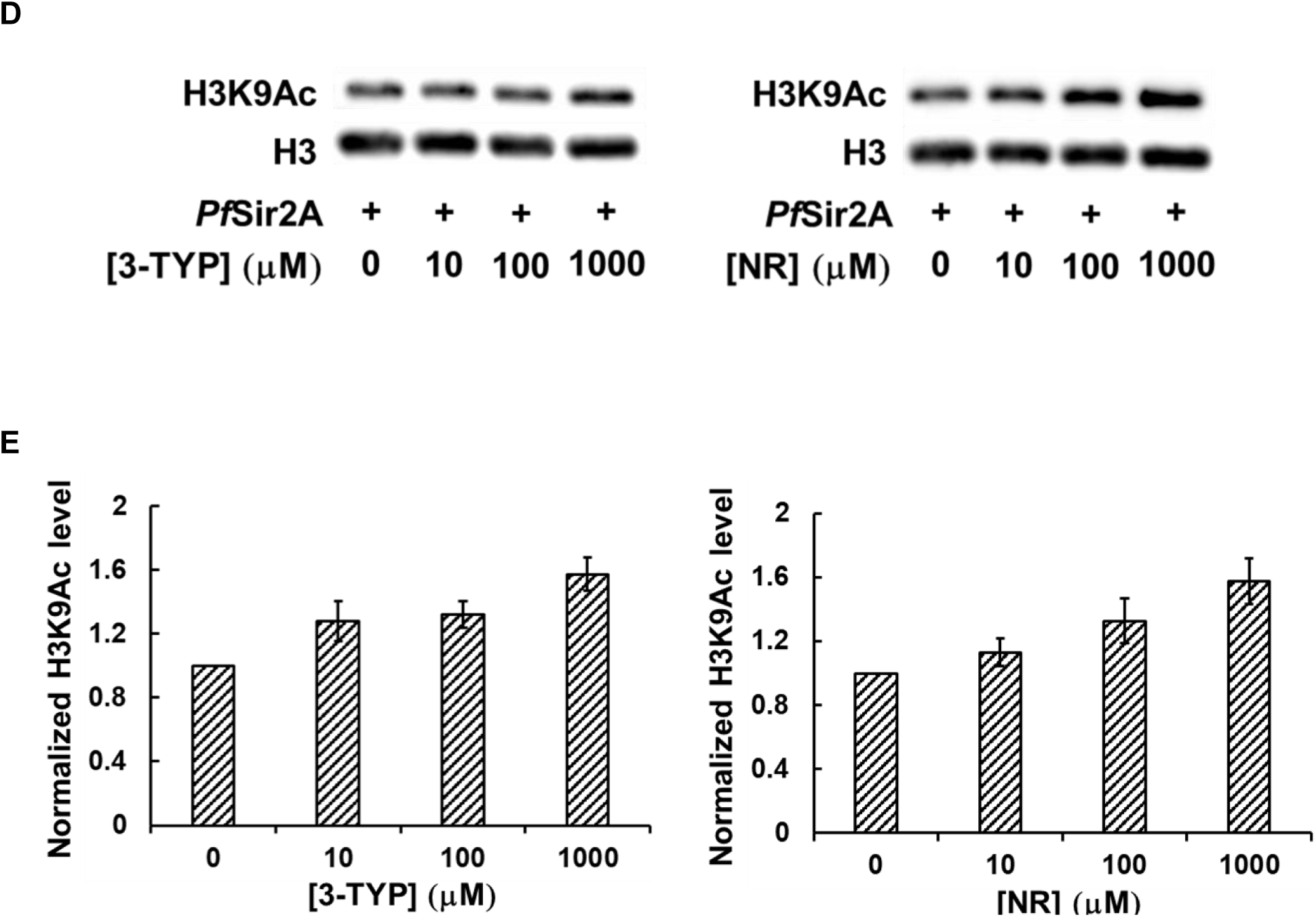
Discovery of *Pf*Sir2A inhibitor from a human sirtuin regulator library. (A) Screening of human sirtuin regulators (from Selleckchem) on *Pf*Sir2A-catalyzed peptide deacetylation at 10, 100, and 1000 µM concentrations; (B)(C) Concentration-dependent inhibition of *Pf*Sir2A-mediated deacetylation by NR (B) or 3-TYP (C); (D) Representative western blots showing that 3-TYP and NR inhibit *Pf*Sir2A deacetylation of H3K9Ac in calf thymus histone; (E) Quantification of the western blot results.

3-TYP is a SIRT3-selective inhibitor with an IC_50_ value of 38 ± 5 µM.^32^ This compound also exhibits cellular activity: incubation of the cells with 3-TYP leads to increased mitochondrial acetylation, significant reduced ATP levels, and markedly increased superoxide, consistent with the phenotypes observed in *sirt3*-deficient mice.^32^ Due to its structural similarity to nicotinamide (NAM), it is proposed that 3-TYP and NAM inhibit sirtuins *via* the same mode of action. Indeed, increasing 3-TYP concentration caused significant reduction of the *V*_max_ value, but negligible changes to the *K*_m_ (**Figure S1**), suggesting that this compound is non-competitive with acetyllysine peptide substrate.

NR is a recently discovered NAD^+^ precursor.^33^ It is metabolically transformed by NR kinases (NRK1 and NRK2) to nicotinamide mononucleotide (NMN).^33–35^ Subsequently, NMN can be incorporated into NAD^+^ *via* the Preiss-Handler pathway.^36^ We and others have demonstrated that NR treatment increases cellular NAD^+^ concentrations significantly.^37–39^ And sirtuin activity is known to be activated with elevated intracellular NAD^+^ contents.^40, 41^ Most recently, our group also discovered that NR is a direct activator of human SIRT5.^42^ It selectively activates SIRT5 deacetylation on both the synthetic peptide and physiological substrates.^42^ In the current study, NR is identified as a non-competitive *Pf*Sir2A inhibitor, as evidenced by the Michaelis-Menten kinetic analysis (**Figure S2**).

3-TYP and NR not only inhibit *Pf*Sir2A deacetylation of synthetic peptides, but also suppress its deacetylation of histone proteins. As shown in **Figures 5D** and **5E**, 3-TYP or NR treatment increases the acetylation level of H3K9 in calf thymus histone in a concentration-dependent manner. At the highest concentration tested, both compounds restore the level of H3K9Ac as compared to the no enzyme controls. The structures of 3-TYP and NR will be further optimized for improved potency and selectivity.

## CONCLUSIONS AND PERSPECTIVES

*P. falciparum* is the causative agent responsible for the most severe form of human malaria. The parasite uses antigenic variation to avoid host antibody recognition, leading to greatly extended periods of infection.^43^ This antigenic variation is mediated by the differential control of a family of *var* genes that encode the surface protein *Pf*EMP1.^44^ Each individual parasite expresses only a single *var* gene at a time. All the other family members are maintained in a transcriptionally silent state, a process controlled by epigenetic mechanisms.^45^

Recent studies suggest the key role of *P. falciparum* silent information regulator 2A (*Pf*Sir2A) in regulating *var* gene transcription.^3,8^ Enzymatically, *Pf*Sir2A is a NAD^+^-dependent histone deacetylase, catalyzing the removal of acetyl groups from histone tails.^14,46^ *Pf*Sir2A has been shown to associate with silenced *var* genes and localizes to chromosome-end clusters and telomere repeats within the nucleus.^3^ Furthermore, genetic ablation of *Pf*Sir2A results in the de-repression of a subset of *var* genes.^7^ All of this evidence points to the idea that heterochromatic silencing mediated by *Pf*Sir2A could directly control the expression of *var* virulence genes. *Pf*Sir2A thus serves as a novel therapeutic target for the development of antimalarial drugs.

The weak *in vitro* deacetylase activity of *Pf*Sir2A makes the characterization of its biochemical properties and the development of selective inhibitors extremely challenging. We hypothesize, with strong preliminary data, *Pf*Sir2A is a highly active histone deacetylase, and this activity is nucleosome-dependent. We use a combination of biochemical, chemical biology, and biophysical approaches to uncover the underlying principles leading to the assembly of *Pf*Sir2A-DNA or *Pf*Sir2A-nucleosome complex, and how this binding is coupled to efficient histone deacetylation.

In addition to DNA and nucleosome, cellular FFAs have also been identified as endogenous *Pf*Sir2A regulators. Myristic acid, a C_14_ fatty acid, is able to stimulate the deacetylase activity, but suppress the defatty-acylase activity. The binding of myristic acid in the large, hydrophobic binding pocket of *Pf*Sir2A causes a conformational rearrangement for enhanced deacetylation. The same binding event also prevents fatty-acylated lysine substrate from binding, leading to the inhibition of defatty-acylation. The data are consistent with the hypothesis that the parasite can sense nutrient availability and respond with antigenic switching.

The biological relevance of *Pf*Sir2A inspires us to conduct inhibitor repurposing for the development of *Pf*Sir2A inhibitors. Screening of a commercial library of human sirtuin regulators led to the discovery of two lead compounds: 3-TYP and NR. Both compounds inhibit the deacetylase activity of *Pf*Sir2A with synthetic peptide substrates and histone substrates. The mode of inhibition of the two molecules were further analyzed. Based on these two lead compounds, potent and *Pf*Sir2A-selective inhibitors will be developed using a suite of enzyme kinetic analysis, structure-activity relationship study, and structure-based drug design.

The current study reveals new understandings on the regulation of *Pf*Sir2A activity, and new scaffolds for *Pf*Sir2A inhibitor. It also provides proof of concept to gain mechanistic and chemical insights into complex enzyme reactions involved in epigenetic control and protein modifications.

## ACKNOWLEDGEMENTS

This work was supported in part by 1R01GM143176-01A1 from NIH/NIGMS (to Y.C.), and Jeffress Trust Award in Interdisciplinary Research from Thomas F. and Kate Miller Jeffress Memorial Trust and Bank of America (to Y.C.). MS analysis reported in this manuscript was performed at the Vermont Biomedical Research Network (VBRN) proteomics facility supported by P20GM103449 (NIH/NIGMS).

## REFERENCES

1. Rieke, B., Overview of the worldwide Handling with the Disease in Prevention, Diagnostics and Therapy WHO World Malaria Report 2017. Flugmedizin Tropenmedizin Reisemedizin 2018, 25 (1), 5–5.

2. Cui, L.; Miao, J., Chromatin-mediated epigenetic regulation in the malaria parasite Plasmodium falciparum. Eukaryot Cell 2010, 9 (8), 1138–49.

3. A. Tonkin, C. J.; Carret, C. K.; Duraisingh, M. T.; Voss, T. S.; Ralph, S. A.; Hommel, M.; Duffy, M. F.; Silva, L. M.; Scherf, A.; Ivens, A.; Speed, T. P.; Beeson, J. G.; Cowman, A. F., Sir2 paralogues cooperate to regulate virulence genes and antigenic variation in Plasmodium falciparum. PLoS Biol 2009, 7 (4), e84.

4. Merrick, C. J.; Dzikowski, R.; Imamura, H.; Chuang, J.; Deitsch, K.; Duraisingh, M. T., The effect of Plasmodium falciparum Sir2a histone deacetylase on clonal and longitudinal variation in expression of the var family of virulence genes. Int J Parasitol 2010, 40 (1), 35–43.

5. Swamy, L.; Amulic, B.; Deitsch, K. W., Plasmodium falciparum var gene silencing is determined by cis DNA elements that form stable and heritable interactions. Eukaryot Cell 2011, 10 (4), 530–9.

6. Rivero, F. D.; Saura, A.; Prucca, C. G.; Carranza, P. G.; Torri, A.; Lujan, H. D., Disruption of antigenic variation is crucial for effective parasite vaccine. Nat Med 2010, 16 (5), 551–7, 1p following 557.

7. Freitas-Junior, L. H.; Hernandez-Rivas, R.; Ralph, S. A.; Montiel-Condado, D.; Ruvalcaba-Salazar, O. K.; Rojas-Meza, A. P.; Mancio-Silva, L.; Leal-Silvestre, R. J.; Gontijo, A. M.; Shorte, S.; Scherf, A., Telomeric heterochromatin propagation and histone acetylation control mutually exclusive expression of antigenic variation genes in malaria parasites. Cell 2005, 121 (1), 25–36.

8. Duraisingh, M. T.; Voss, T. S.; Marty, A. J.; Duffy, M. F.; Good, R. T.; Thompson, J. K.; Freitas-Junior, L. H.; Scherf, A.; Crabb, B. S.; Cowman, A. F., Heterochromatin silencing and locus repositioning linked to regulation of virulence genes in Plasmodium falciparum. Cell 2005, 121 (1), 13–24.

9. Scherf, A.; Lopez-Rubio, J. J.; Riviere, L., Antigenic variation in Plasmodium falciparum. Annu Rev Microbiol 2008, 62, 445–70.

10. Mancio-Silva, L.; Rojas-Meza, A. P.; Vargas, M.; Scherf, A.; Hernandez-Rivas, R., Differential association of Orc1 and Sir2 proteins to telomeric domains in Plasmodium falciparum. J Cell Sci 2008, 121 (Pt 12), 2046–53.

11. Lopez-Rubio, J. J.; Mancio-Silva, L.; Scherf, A., Genome-wide analysis of heterochromatin associates clonally variant gene regulation with perinuclear repressive centers in malaria parasites. Cell Host Microbe 2009, 5 (2), 179–90.

12. Mancio-Silva, L.; Lopez-Rubio, J. J.; Claes, A.; Scherf, A., Sir2a regulates rDNA transcription and multiplication rate in the human malaria parasite Plasmodium falciparum. Nat Commun 2013, 4, 1530.

13. Merrick, C. J.; Duraisingh, M. T., Plasmodium falciparum Sir2: an unusual sirtuin with dual histone deacetylase and ADP-ribosyltransferase activity. Eukaryot Cell 2007, 6 (11), 2081–91.

14. French, J. B.; Cen, Y.; Sauve, A. A., Plasmodium falciparum Sir2 is an NAD+-dependent deacetylase and an acetyllysine-dependent and acetyllysine-independent NAD+ glycohydrolase. Biochemistry 2008, 47 (38), 10227–39.

15. Banerjee, C.; Nag, S.; Goyal, M.; Saha, D.; Siddiqui, A. A.; Mazumder, S.; Debsharma, S.; Pramanik, S.; Bandyopadhyay, U., Nuclease activity of Plasmodium falciparum Alba family protein PfAlba3. Cell Rep 2023, 42 (4), 112292.

16. Zhu, A. Y.; Zhou, Y.; Khan, S.; Deitsch, K. W.; Hao, Q.; Lin, H., Plasmodium falciparum Sir2A preferentially hydrolyzes medium and long chain fatty acyl lysine. ACS Chem Biol 2012, 7 (1), 155–9.

17. Dalmasso, M. C.; Carmona, S. J.; Angel, S. O.; Aguero, F., Characterization of Toxoplasma gondii subtelomeric-like regions: identification of a long-range compositional bias that is also associated with gene-poor regions. BMC Genomics 2014, 15 (1), 21.

18. Lowary, P. T.; Widom, J., New DNA sequence rules for high affinity binding to histone octamer and sequence-directed nucleosome positioning. J Mol Biol 1998, 276 (1), 19–42.

19. Hoff, K. G.; Avalos, J. L.; Sens, K.; Wolberger, C., Insights into the sirtuin mechanism from ternary complexes containing NAD+ and acetylated peptide. Structure 2006, 14 (8), 1231–40.

20. Fameau, A. L.; Ventureira, J.; Novales, B.; Douliez, J. P., Foaming and emulsifying properties of fatty acids neutralized by tetrabutylammonium hydroxide. Colloids and Surfaces a-Physicochemical and Engineering Aspects 2012, 403, 87–95.

21. Tholstrup, T.; Sandstrom, B.; Bysted, A.; Holmer, G., Effect of 6 dietary fatty acids on the postprandial lipid profile, plasma fatty acids, lipoprotein lipase, and cholesterol ester transfer activities in healthy young men. Am J Clin Nutr 2001, 73 (2), 198–208.

22. Xu, J.; Christian, B.; Jump, D. B., Regulation of rat hepatic L-pyruvate kinase promoter composition and activity by glucose, n-3 polyunsaturated fatty acids, and peroxisome proliferator-activated receptor-alpha agonist. J Biol Chem 2006, 281 (27), 18351–62.

23. Jump, D. B.; Clarke, S. D.; Thelen, A.; Liimatta, M., Coordinate regulation of glycolytic and lipogenic gene expression by polyunsaturated fatty acids. J Lipid Res 1994, 35 (6), 1076–84.

24. Lahree, A.; Mello-Vieira, J.; Mota, M. M., The nutrient games - Plasmodium metabolism during hepatic development. Trends Parasitol 2023, 39 (6), 445–460.

25. Amiar, S.; Katris, N. J.; Berry, L.; Dass, S.; Duley, S.; Arnold, C. S.; Shears, M. J.; Brunet, C.; Touquet, B.; McFadden, G. I.; Yamaryo-Botte, Y.; Botte, C. Y., Division and Adaptation to Host Environment of Apicomplexan Parasites Depend on Apicoplast Lipid Metabolic Plasticity and Host Organelle Remodeling. Cell Rep 2020, 30 (11), 3778–3792 e9.

26. Schneider, V. M.; Visone, J. E.; Harris, C. T.; Florini, F.; Hadjimichael, E.; Zhang, X.; Gross, M. R.; Rhee, K. Y.; Ben Mamoun, C.; Kafsack, B. F. C.; Deitsch, K. W., The human malaria parasite Plasmodium falciparum can sense environmental changes and respond by antigenic switching. Proc Natl Acad Sci U S A 2023, 120 (17), e2302152120.

27. Sauve, A. A.; Schramm, V. L., Sir2 regulation by nicotinamide results from switching between base exchange and deacetylation chemistry. Biochemistry 2003, 42 (31), 9249–56.

28. Avalos, J. L.; Bever, K. M.; Wolberger, C., Mechanism of sirtuin inhibition by nicotinamide: altering the NAD(+) cosubstrate specificity of a Sir2 enzyme. Mol Cell 2005, 17 (6), 855–68.

29. Prusty, D.; Mehra, P.; Srivastava, S.; Shivange, A. V.; Gupta, A.; Roy, N.; Dhar, S. K., Nicotinamide inhibits Plasmodium falciparum Sir2 activity in vitro and parasite growth. FEMS Microbiol Lett 2008, 282 (2), 266–72.

30. Chakrabarty, S. P.; Ramapanicker, R.; Mishra, R.; Chandrasekaran, S.; Balaram, H., Development and characterization of lysine based tripeptide analogues as inhibitors of Sir2 activity. Bioorg Med Chem 2009, 17 (23), 8060–72.

31. Curry, A. M.; White, D. S.; Donu, D.; Cen, Y., Human Sirtuin Regulators: The “Success” Stories. Front Physiol 2021, 12, 752117.

32. Galli, U.; Mesenzani, O.; Coppo, C.; Sorba, G.; Canonico, P. L.; Tron, G. C.; Genazzani, A. A., Identification of a sirtuin 3 inhibitor that displays selectivity over sirtuin 1 and 2. Eur J Med Chem 2012, 55, 58–66.

33. Tempel, W.; Rabeh, W. M.; Bogan, K. L.; Belenky, P.; Wojcik, M.; Seidle, H. F.; Nedyalkova, L.; Yang, T.; Sauve, A. A.; Park, H. W.; Brenner, C., Nicotinamide riboside kinase structures reveal new pathways to NAD+. PLoS Biol 2007, 5 (10), e263.

34. Bogan, K. L.; Brenner, C., Nicotinic acid, nicotinamide, and nicotinamide riboside: a molecular evaluation of NAD+ precursor vitamins in human nutrition. Annu Rev Nutr 2008, 28, 115–30.

35. Fletcher, R. S.; Ratajczak, J.; Doig, C. L.; Oakey, L. A.; Callingham, R.; Da Silva Xavier, G.; Garten, A.; Elhassan, Y. S.; Redpath, P.; Migaud, M. E.; Philp, A.; Brenner, C.; Canto, C.; Lavery, G. G., Nicotinamide riboside kinases display redundancy in mediating nicotinamide mononucleotide and nicotinamide riboside metabolism in skeletal muscle cells. Mol Metab 2017, 6 (8), 819–832.

36. Preiss, J.; Handler, P., Biosynthesis of diphosphopyridine nucleotide. I. Identification of intermediates. J Biol Chem 1958, 233 (2), 488–92.

37. Tran, A.; Yokose, R.; Cen, Y., Chemo-enzymatic synthesis of isotopically labeled nicotinamide riboside. Org Biomol Chem 2018, 16 (19), 3662–3671.

38. Trammell, S. A.; Schmidt, M. S.; Weidemann, B. J.; Redpath, P.; Jaksch, F.; Dellinger, R. W.; Li, Z.; Abel, E. D.; Migaud, M. E.; Brenner, C., Nicotinamide riboside is uniquely and orally bioavailable in mice and humans. Nat Commun 2016, 7, 12948.

39. Martens, C. R.; Denman, B. A.; Mazzo, M. R.; Armstrong, M. L.; Reisdorph, N.; McQueen, M. B.; Chonchol, M.; Seals, D. R., Chronic nicotinamide riboside supplementation is well-tolerated and elevates NAD(+) in healthy middle-aged and older adults. Nat Commun 2018, 9 (1), 1286.

40. Nakagawa, T.; Lomb, D. J.; Haigis, M. C.; Guarente, L., SIRT5 Deacetylates carbamoyl phosphate synthetase 1 and regulates the urea cycle. Cell 2009, 137 (3), 560–70.

41. Belenky, P.; Racette, F. G.; Bogan, K. L.; McClure, J. M.; Smith, J. S.; Brenner, C., Nicotinamide riboside promotes Sir2 silencing and extends lifespan via Nrk and Urh1/Pnp1/Meu1 pathways to NAD+. Cell 2007, 129 (3), 473–84.

42. Curry, A. M.; Rymarchyk, S.; Herrington, N. B.; Donu, D.; Kellogg, G. E.; Cen, Y., Nicotinamide riboside activates SIRT5 deacetylation. FEBS J 2023, 290 (19), 4762–4776.

43. Geleta, G.; Ketema, T., Severe Malaria Associated with Plasmodium falciparum and P. vivax among Children in Pawe Hospital, Northwest Ethiopia. Malar Res Treat 2016, 2016, 1240962.

44. Deitsch, K. W.; Dzikowski, R., Variant Gene Expression and Antigenic Variation by Malaria Parasites. Annu Rev Microbiol 2017, 71, 625–641.

45. Cortes, A.; Deitsch, K. W., Malaria Epigenetics. Cold Spring Harb Perspect Med 2017, 7 (7).

46. Chakrabarty, S. P.; Saikumari, Y. K.; Bopanna, M. P.; Balaram, H., Biochemical characterization of Plasmodium falciparum Sir2, a NAD+-dependent deacetylase. Mol Biochem Parasitol 2008, 158 (2), 139–51.

