## Supplementary Figures for "Biochemical Characterization and Inhibitor Discovery for *Pf*Sir2A – New Tricks for An Old Enzyme"

**Figure S1.** Michealis-Menten kinetic analysis of 3-TYP inhibition.

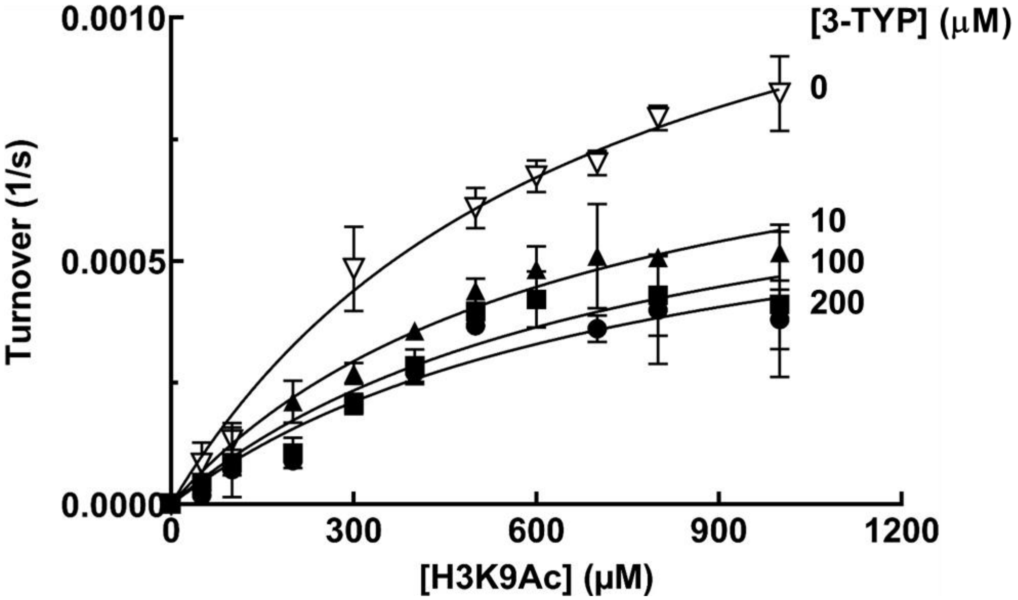


The mode of inhibition analysis of 3-TYP was performed as described in "Methods and Materials". With increasing concentrations of 3-TYP, the $K_m$ value remains roughly unchanged, while the $V_{max}$ is significantly reduced.

**Figure S2.** Michealis-Menten kinetic analysis of NR inhibition.

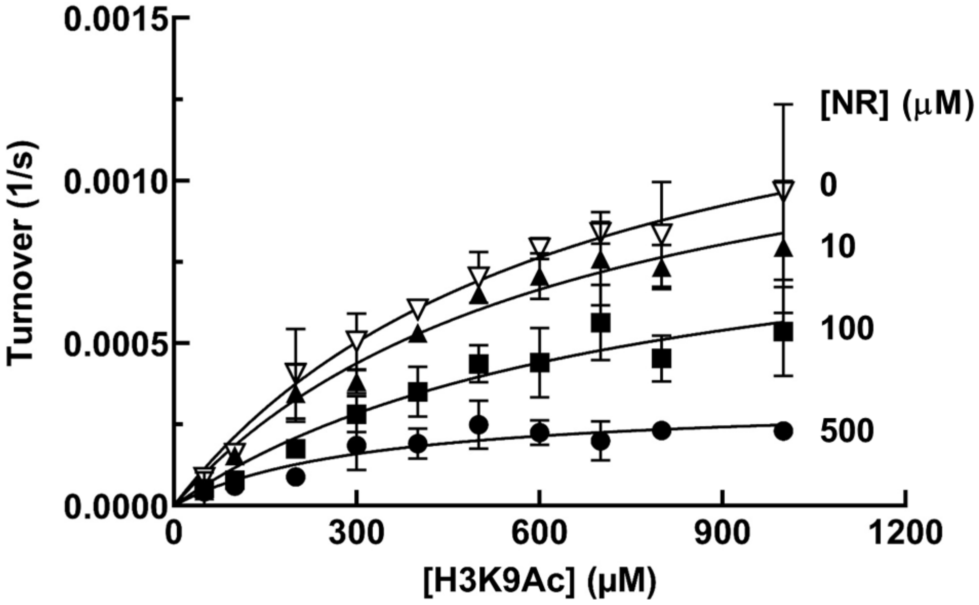


The mode of inhibition analysis of NR was performed as described in “Methods and Materials”. Increasing NR concentration leads to markedly decreased $V_{max}$, but negligible changes to the $K_m$.
